# Strengthening the BioCompute Standard by Crowdsourcing on PrecisionFDA

**DOI:** 10.1101/2020.11.02.365528

**Authors:** Sarah H Stephens, Charles Hadley King, Sean Watford, Janisha Patel, Dennis A. Dean, Soner Koc, Nan Xiao, Eric F. Donaldson, Elaine E. Thompson, Anjan Purkayastha, Raja Mazumder, Elaine Johanson, Jonathon Keeney

## Abstract

**Background:** The field of bioinformatics has grown at such a rapid pace that a gap in standardization exists when reporting an analysis. In response, the BioCompute project was created to standardize the type and method of information communicated when describing a bioinformatic analysis. Once the project became established, its goals shifted to broadening awareness and usage of BioCompute, and soliciting feedback from a larger audience. To address these goals, the BioCompute project collaborated with precisionFDA on a crowdsourced challenge that ran from May 2019 to October 2019. This challenge had a beginner track where participants submitted BCOs based on a pipeline of their choosing, and an advanced track where participants submitted applications supporting the creation of a BCO and verification of BCO conformance to specifications.

**Results:** In total, there were 28 submissions to the beginner track (including submissions from a bioinformatics master’s class at George Washington University) and three submissions to the advanced track. Three top performers were selected from the beginner track, while a single top performer was selected for the advanced track. In the beginner track, top performers differentiated themselves by submitting BCOs that included more than the minimally compliant content. Advanced track submissions were very impressive. They included a complete web application, a command line tool that produced a static result, and a dockerized container that automatically created the BCO as the tool was run. The ability to harmonize the correct function, a simple user experience, and the aesthetics of the tool interface differentiated the tools.

**Conclusions:** Despite being new to the concept, most beginner track scores were high, indicating that most users understood the fundamental concepts of the BCO specification. Novice bioinformatics students were an ideal cohort for this Challenge because of their lack of familiarity with BioCompute, broad diversity of research interests, and motivation to submit high-quality work. This challenge was successful in introducing the BCO to a wider audience, obtaining feedback from that audience, and resulting in a tool novices may use for BCO creation and conformance. In addition, the BCO specification itself was improved based on feedback illustrating the utility of a “wisdom of the crowd” approach to standards development.

## Background

The increasing pace of development in bioinformatic analyses is contributing to rapid progress throughout the life sciences. However, because of the relatively new and burgeoning nature of the bioinformatics field, there is a lack of standardization when reporting a bioinformatic analysis workflow, leading to a breakdown of communication and making it difficult to understand and potentially impossible to reproduce. In response, a community of individuals has formed the BioCompute project [1, 2], meant to standardize the type and method of information communicated when describing a bioinformatic analysis.

The need for what would become the BioCompute project was officially documented at the FDA in 2013 by the Genomics Working Group (GWG). The GWG, which consists of subject matter experts from across the FDA, authored an internal document that outlined the needs of FDA reviewers in evaluating NGS analysis.

The document generated by the GWG formed the framework for what would ultimately become the BioCompute specification by outlining four requirements for the project. Namely, any solution should: 1) be human-readable, 2) be computer readable (i.e., structured fields with a well-defined scope of information), 3) contain enough information to understand the bioinformatics pipelines, interpret information, maintain records, and reproduce experiments, and 4) have some way to make sure the information has not been altered.

Through a series of workshops, input was collected from various interested parties, and an initial specification was generated (https://w3id.org/biocompute/1.0.1). Subsequent input through additional workshops and use case trials was used to refine the specification further, resulting in additional iterations [https://github.com/biocompute-objects/BCO_Specification], and culminating in what is now an IEEE-approved standard (2791-2020)(https://standards.ieee.org/standard/2791-2020.html), which is associated with an open source code repository (https://w3id.org/ieee/ieee-2791-schema/). The IEEE standard is the result of hundreds of participants in academia, regulatory, and private industry settings, who participated in hands-on workshops, conferences, and in two publications[1, 2]. The FDA has since announced support for the use of the IEEE-approved standard 2791-2020 in regulatory submissions; more information can be found on the Federal Register (<u>85 FR 44304</u>).

BioCompute is a human and machine-readable specification for reporting analysis pipelines. A specific instance of a pipeline that adheres to the BioCompute specification is a BioCompute Object (BCO). Fields in a BCO are categorically grouped in an intuitive way into “Domains.” Programmatic domains (Description Domain, Execution Domain, and Parametric Domain) include granular details, like scripts, dependencies, and any other prerequisites required to execute, all of which were organized in a consensus-driven way. BCOs include a hash function for integrity verification, and an embargo field for BCOs to be made public after a certain date.

BioCompute is built on the JSON Schema (http://json-schema.org/draft-07/schema#), and is an amalgam of several important concepts, including 1) a “Usability Domain” for researchers to include plain text descriptions of what was done in their workflow and why, and to describe the scientific question and any other relevant information; 2) data provenance that tracks the flow of data from source to final analysis and 3) a “Verification Kit” that allows a Reviewer to quickly check the validity of results against known error rates in the analysis, and which consists of the IO Domain and the Error Domain. As well as tracking data flow, the data provenance domain includes detailed user attribution (based on Provenance Authoring and Versioning [2]) for both the generator of the BCO and the reviewer.

The Error Domain is comprised of information that demarcates the boundary of a pipeline, such as minimum depth of coverage, and statistical descriptors, like false positive/negative rates. The Verification Kit is a way of combining the IO Domain and the Error Domain, such that one can evaluate the validity of a set of output files, given a set of input files and the error in the pipeline. Because the BCO project is maintained as an open source repository, creative, non-standard uses can be branched, such as an internal schema that organizes proprietary data not visible to the public, but which is still compatible with the standard.

A workshop hosted at the FDA in May of 2019 further articulated the need and refined the scope of BioCompute, showcased use case examples, demonstrated tools that made use of the BioCompute specification, and presented an update on the progress of publishing the IEEE standard. It was clear that the BioCompute project was established and had great momentum, and so the goal has largely shifted to broadening awareness and usage of BioCompute and soliciting feedback from a larger audience. Prior to the challenge, this larger audience consisted of 1) those completely unfamiliar with the project and 2) bioinformatics novices.

Crowdsourcing presented a good opportunity to leverage the benefits of a larger audience, engage new stakeholders, and generate a diversity of insights into the evolving standard. Citizen science efforts such as crowdsourcing are well established methods to generate benefits over internal ideation including generating unexpected solutions to problems. Competition platforms like Innocentive, Kaggle, and DREAM offer sponsors a one-stop shop to organize, run, and evaluate results for their challenge. PrecisionFDA is one such community-based platform that leverages the benefit of a crowdsourcing model to host challenges aimed at developing innovations to inform the science of precision medicine and the advancement of regulatory science. What precisionFDA offers beyond a competition platform is a high-performance computing environment where users can host, analyze, and collaborate with experts in a secure (FEDRAMP and FISMA moderate) space. Since precisionFDA is a community-based platform and not just a challenge platform, it provides collaborators with both the ability to directly engage with the regulatory and bioinformatics community and a unique opportunity to collaborate on novel tools or specification development and application. These features make precisionFDA a natural mechanism for engaging the bioinformatics community.

The BioCompute project and precisionFDA collaborated on a crowdsourced challenge that ran from May 2019 to October 2019 with the goals of 1) broadening awareness and usage of BioCompute, 2) engaging a larger “test” audience for the standard, 3) soliciting feedback from a larger audience, and 4) lowering the barrier to entry for those unfamiliar with standards compliance or some other aspect of BCOs. The challenge had a beginner track where participants submitted BCOs based on a pipeline of their choosing, and an advanced track where participants submitted applications supporting creation of a BCO and conformance to specifications. This article presents challenge results, achievements, and conclusions.

Figure 1 shows a schematic diagram of the workflows for both beginner and advanced track submissions to this challenge

**Figure 1.**
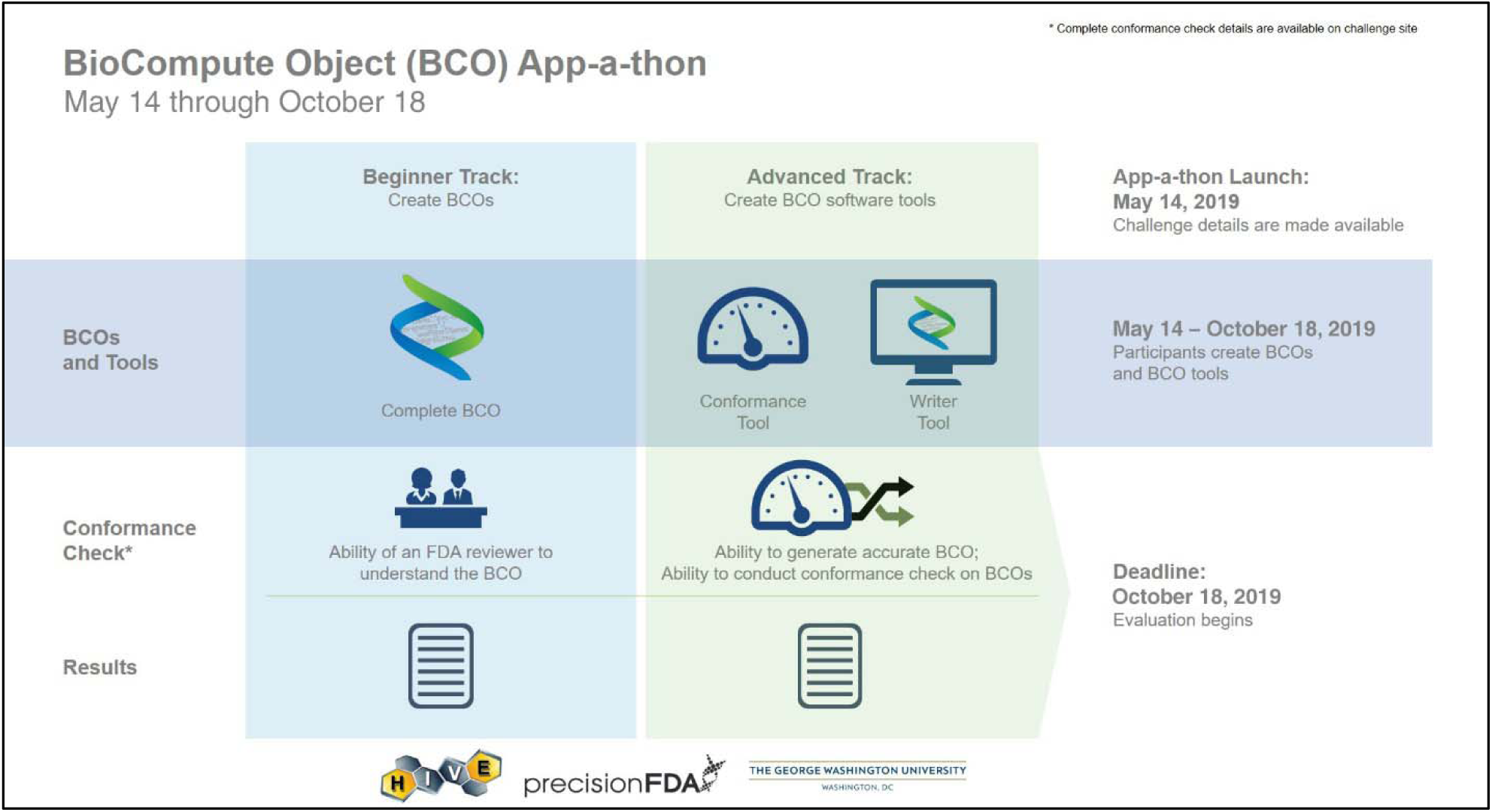
Challenge Workflows.

## Methods

### Beginner Track

To encourage novice users to participate, users were encouraged to use one of two BCO writer/editor tools already developed by the community: a standalone form-based editor on the web or a tool integrated into the Cancer Genomics Cloud (CGC). Participants in the beginner track were asked to create a BCO that documented a multi-step bioinformatics pipeline, conforming to the BioCompute specification that was current at the time. Additionally, submissions were to be in JSON file format and not contain proprietary or sensitive information as they would be publicly available post-challenge.

### The Stand-Alone Editor Tool

The BCO Editor is a user-friendly tool designed to make BioCompute accessible to researchers with minimal technical knowledge of JSON or BioCompute. Challenge participants with no prior knowledge of JSON or BioCompute were able to use the tool to create, modify, and download BCOs for submission to the challenge. To accommodate varying technical skill levels of users, the tool provides the option of either using a form-based format or manually editing the raw JSON. In addition to an example BCO made accessible on the tool, all BCO fields are accompanied with field descriptions to familiarize users with BCO concepts as they are developing their BCO. Submitted BCOs are validated against the current BioCompute JSON schema and stored on the platform. For proprietary or unpublished research, an embargo period can be specified in the BCO at the user’s discretion, after which the BCO will be publicly accessible for viewing and downloading. Challenge participants who used the Editor were encouraged to submit any issues they encountered to the BCO Editor GitHub repository, allowing the development team to troubleshoot user issues and receive valuable UX feedback in real-time. While the legacy version of the Editor used during the challenge is depreciated, objects created during this challenge have been migrated to the new BCO Portal at https://portal.aws.biochemistry.gwu.edu/ where all current and future releases will be hosted.

### Advanced Track

This challenge offered an advanced track in the form of an app-a-thon (application competition) where participants submitted applications supporting the creation and display of a BCO, the ability to check conformance against the BioCompute specification, and a user manual (i.e., README explanation of intended use, installation instructions, and other pertinent info). Participants could submit in one of two ways: direct upload of a compiled app to the precisionFDA site, or source code submission. In addition, the challenge allowed for an application to be submitted that was fully integrated into an existing platform as long as the application was accessible to the evaluation team (e.g., a URL to a page where the application can be tested).

### Student recruitment

Bioinformatics masters students in the BIOC 6223 Introduction to Bioinformatics course at George Washington University were recruited to participate in the challenge as part of the course curriculum. Students were introduced to the core concepts of the JSON Schema (http://json-schema.org/draft-07/schema#), basics of analysis pipelines, database usage, and BioCompute in the first half of the semester. Additionally, the Challenge was incorporated into the midterm. Students were required to research a cancer-related bioinformatics pipeline that made use of open source repositories, and to build a BCO representation of that pipeline through a three and a half week process. Students in the course had little to no bioinformatics experience and represented an ideal test audience for the BioCompute standard.

### Grading criteria

#### Beginner Track

Each submission was distributed to one of four reviewers, who evaluated the BCO within three sections: basic qualifications, documentation, and bonus. Basic qualifications for a BCO included provenance, prerequisites, and scripts (or working pointers). In addition, the BCO was required to document a valid computational pipeline and conform with specifications (1.3.0) based on the validate.py script (https://w3id.org/biocompute/1.3.1/validate.py). The documentation score was based on the inclusion of a detailed Usability Domain with the purpose of the pipelines, references to help understand the application of the pipelines, and links to any associated materials that would help understand application of the pipeline. Documentation also included a clearly named Description Domain and documented pipeline steps. Bonus points were awarded for creative use of keywords, quality of the Usability Domain, inclusion of optional domains (Extension and Error Domains), and reproducibility of the BCO. Scoring was in keeping with the spirit of the BCO standard, to communicate a workflow in a way that is easy to understand. Bonus scoring for “creative use” encompassed the way in which the submitted domains worked together coherently and concisely to tell the story of the multi-step workflow.

#### Advanced Track

Submissions to the advanced track were evaluated within five sections: basic qualifications, functionality, documentation, usability and aesthetics, and bonus. Basic qualifications included the ability of the application to create, display, and check conformance of a BCO to BioCompute Specification 1.3.0, as well as a verified digital signature. For a tool to be functional, it had to take user input to generate a BCO, correctly identify and report deviations or errors from specifications, and display a BCO in a user-friendly interface. The documentation score was based on the inclusion of a plain text README, including the purpose of the application, references to help understand the application, links to associated materials that would help understand the application, and clearly named and documented steps to create a BCO. Usability and aesthetics included installation instructions with a working example, interface design and ease of user interactions, user-friendly conformance checking, and tooltips, and prompts to help users with fields to be filled out manually. Bonus points were awarded for applications that could easily update when new versions of the standard are released, handle more than one version of the specification, compare more than one BCO at a time, and support following links and URLs in a BCO.

## Results

### Overall

In total, there were 28 submissions to the beginner track and 3 submissions to the advanced track (**Table 1**). Three top performers were selected from the beginner track, while a single top performer was selected for the advanced track based on the evaluation criteria.

**Table 1.** Summary of Results

|  | Basic Qualifications | Documentation | Bonus | Total |
| --- | --- | --- | --- | --- |
| Number of Submissions | 28 | 28 | 28 | 28 |
| Mean Score | 22.32 | 4.32 | 3.75 | 30.4 |
| Standard Deviation | 3.96 | 0.94 | 2.01 | 5.12 |
| Lowest Score | 15 | 2 | 1 | 20 |
| Highest Score | 25 | 5 | 8 | 38 |
| Total Possible Points | 25 | 5 | 10 | 40 |

### Beginner Track

BCO submissions were either based on the participants own workflow, or derived from a workflow from a published manuscript (3-9). Top performers were selected based on total score. **Table 2** names the top three performers, and shows their overall score as well as their scores within each section.

**Table 2.** Challenge Top Performers

|  | Score total<br>(out of 40) | Basic Qualifications Score<br>(out of 25) | Documentation Score<br>(out of 5) | Bonus Score<br>(out of 10) |
| --- | --- | --- | --- | --- |
| Najuma Babirye | 38 | 25 | 5 | 8 |
| Elizabeth Lee | 38 | 25 | 5 | 8 |
| Nirmala Ramnarine | 37 | 25 | 5 | 7 |

Figure displays score trends for each section. A normalized score was calculated by dividing the score of each submission by the total possible points from each section (including total points). The basic qualifications and documentation sections were summed and capture a majority of the total possible points (30 points or 75% of all possible points). Figure 2(A) shows anonymized scores for each participant. In general, submissions performed well in both basic qualifications and documentation sections and the majority had combined, normalized scores over 80% (B). Bonus scores were lower and more broadly distributed (C), leading to a broader distribution of overall scores (D). The scores for the combined basic qualifications and documentation sections scores, which were required for the midterms of student participants, failed to show a correlation with the scores on the bonus section (r=0.4, p=0.04) which was only part of the Challenge. Overall, the top performing submissions scored high in all three sections, but the bonus section set top performers apart from the rest of the submissions.

**Figure 2.**
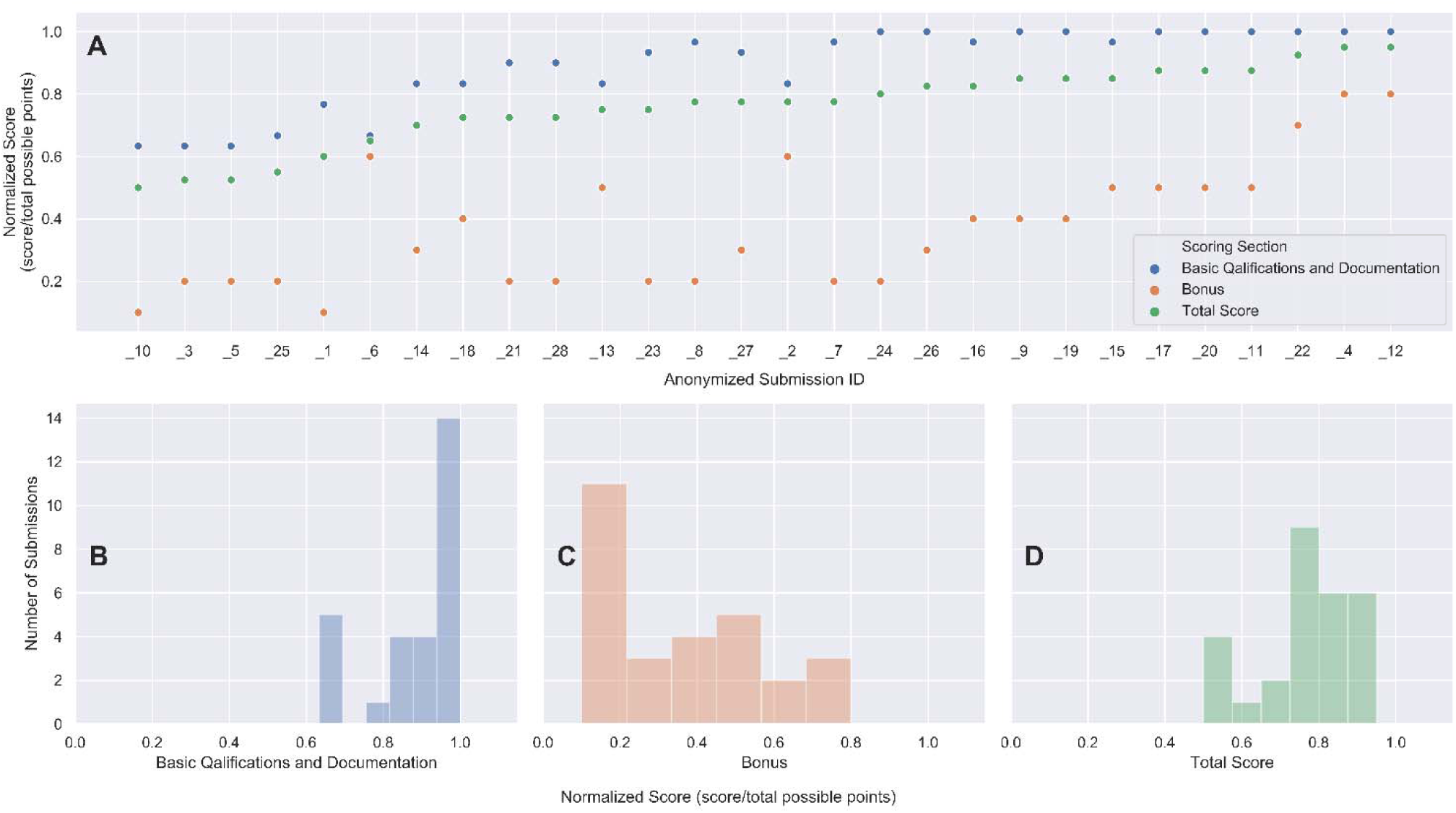
Score trends across each section of the evaluation criteria.

Top performers differentiated themselves by submitting BCOs that included more than the minimally compliant content. For example, an average submission linked the paper(s) from which the pipeline was taken while top performers linked to the paper as well as other resources (such as through “xref” fields) aimed to help a reader develop a deeper understanding of concepts relevant to the paper or pipeline. Additionally, six of the 28 submissions included either an error or extension domain whereas top performers were the only participants to include both.

In addition to awarding top performers, participants of the beginner track of this challenge were awarded badges based on additional categories. For the categories Functional, Documentation, and Usability, participants could receive bronze, silver, or gold badges based on their score. For the all-or-nothing categories of “All the Basics” and “Extra Credit,” participants could receive a platinum badge. A list of participant badges is shown below in Figure 3.

**Figure 3.**
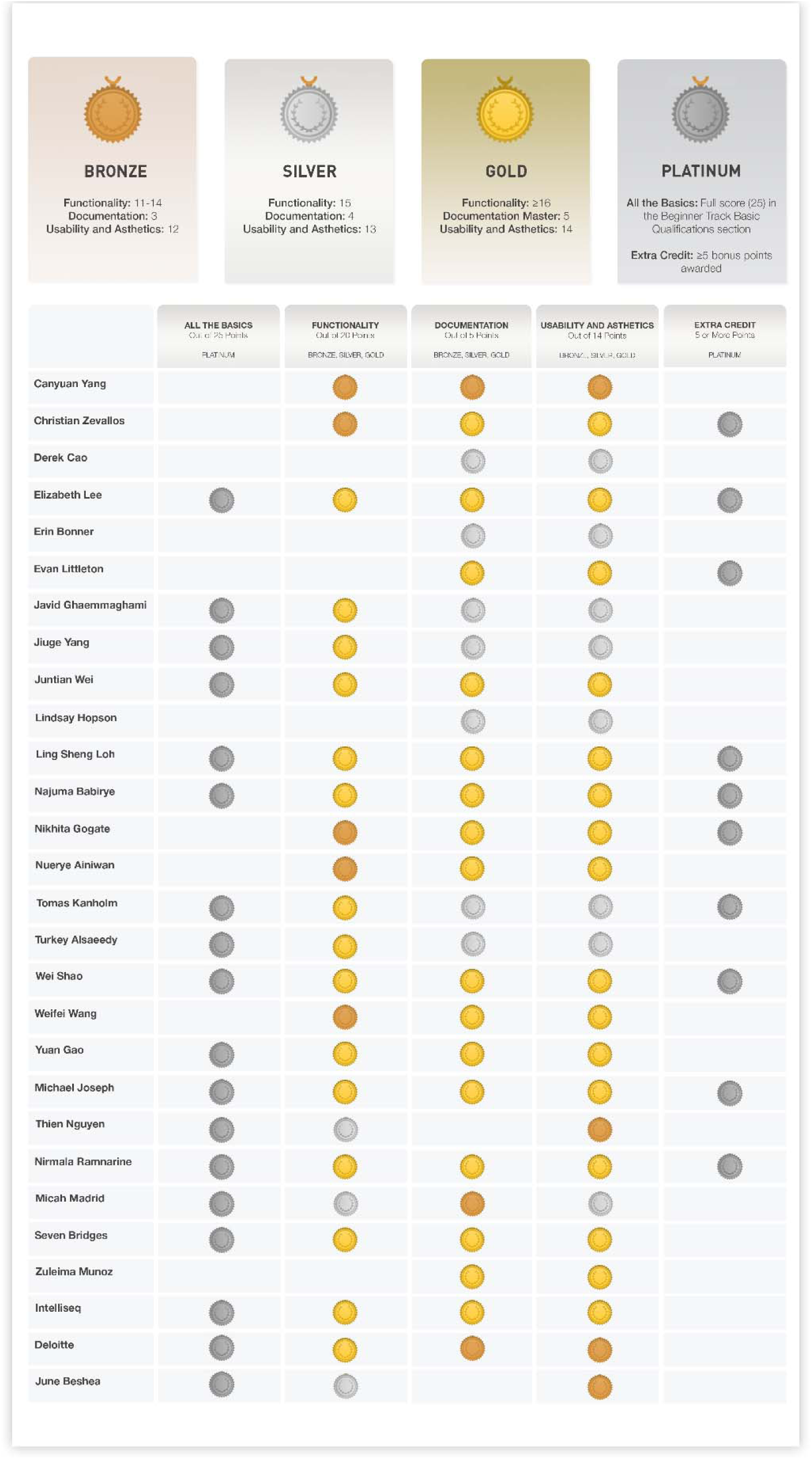
Beginner track participant badges.

### Advanced Track

A single top performing team was selected for the advanced track based on the evaluation criteria: Seven Bridges Genomics team consisting of Drs Nan Xiao and Dennis Dean, and Soner Koc [10].

Overall, there was a great deal of variability in the function and creativity of the three advanced track submissions. Advanced track submissions were very impressive. They included a complete web application, a command line tool that produced a static result, and a dockerized container that automatically created the BCO as the tool was run. The goal was to develop applications that lower the barrier to entry for those unfamiliar with standards compliance, JSON, or some other aspect of BCOs. The instructions and basic requirements provided to the applicants provided the desired final outcome but did not specify how those outcomes should be achieved. The ability to harmonize the correct function, a simple user experience, and the aesthetics of the tool interface (hinted at in the instructions, but not explicitly required) differentiated the tools. Each of the submitted tools was markedly different in how documentation was employed, execution environment utilized, and in the overall aesthetics.

When evaluating, reviewers put emphasis on the ability to install and run the application. All three submissions provided great detail on important analysis pipelines, but none contained parametric domain data. Description of parameters is among the most important criteria for an actual regulatory submission. However, because the Parametric Domain is only required for any parameter that is changed from the default and therefore not required broadly (in the event that every tool in a pipeline is used in a “canned” way without changing any parameters), it is currently not possible to know if these were omitted accidentally. Best practices guidelines have therefore been revised to suggest that a BCO generator indicates in the Usability Domain when a Parametric Domain is intentionally left blank.

## Discussion

Despite being new to the BCO concept, most beginner track scores were high, indicating that most users understood the fundamental concepts of BCO specification. The high combined scores on basic qualification and documentation and the lack of correlation between core and bonus scores suggests that many participants were focused on the core content. This is not surprising because for student participants, the incorporation of additional domains beyond those required in the specification was not required for midterm scoring criteria. Participants submitting optional domains or details aimed at earning bonus points may have been more interested in achieving high scores in the Challenge. Lack of inclusion of optional material in the Challenge can therefore only be considered qualitatively.

Novice bioinformatics students were an ideal cohort for this Challenge because of their lack of familiarity with BioCompute, broad diversity of research interests, and motivation to submit high quality work through Challenge incorporation into the course curriculum in the form of a midterm. Students also posed many questions and thoroughly explored the idea space.

Expanded use of BCOs may come with more complex and nuanced pipelines, requiring more in-depth documentation, and therefore an expanded use of additional elements. In particular, the Error Domain may be important for explicitly demarcating the detection abilities of a pipeline or false negative rates. Because every computational analysis is different, the contents of the Error Domain for each analysis will be different. The Error Domain was therefore determined to be too difficult to standardize at this time and thus was not included as a required domain in the current IEEE 2791-2020 standard. However, it is strongly recommended through field tests such as the one described here that the Error Domain be included, as it can provide the needed context for understanding results (“do the results in the output fall within the level of detection in the Error Domain?”) or reproducing the analysis (“do I have enough depth of coverage to be able to run this pipeline?”).

Additionally, the Extension Domain may see expanded use. The BioCompute project was developed through a consensus approach among a large community of bioinformatics experts, and the base BCO schema is therefore expected to meet most needs. However, the Extension Domain was an ideal place for Challenge participants who wanted to represent other information that was not critical to the analysis, but which they felt was relevant (e.g., microarray data). Following the Challenge, the Extension Domain was included in 2791-2020 as an optional domain that must be in the form of a schema, against which a BCO can be validated for conformance. Examples of Extension Domains have also been generated[https://biocomputeobject.org/BCOCheatSheet.pdf].

Students were a good “experimental cohort” and test audience because they provided a great diversity of use cases with varying types of pipelines and were heavily incentivized to submit high-quality work, since it was their entire midterm grade (and publicly viewable). Typical volunteer participants cannot be incentivized in the same way. An industry sponsor in a real-life setting may also be unfamiliar with the standard, but would be incentivized to create a BCO to the best of their ability since it could be a useful addition to a formal submission to the FDA. Industry interest in the BCO is indicated by the participation of industry in the advanced track. The advanced track had a much higher barrier to entry, yet three submissions, all from industry (Seven Bridges Genomics, Deloitte, and Intelliseq), were received.

Overall, this challenge was successful in that it introduced the concept to a wider audience, obtained feedback from that audience, and resulted in a tool that novices can use for BCO creation and conformance. The results from this challenge provides insights for future app-a-thons to have a more refined set of “best practices” (or outright guidelines) regarding the construction of tools for submission, while still leaving room for creative interpretation. In addition, this challenge achieved a measurable stride in lowering the barrier to implementation of the BioCompute standard by using participant feedback to improve the specification. The following paragraphs detail additional tools that were not utilized for the challenge but will be useful to those looking to incorporate the specification into their published workflow.

### The HIVE-integrated tool

In order for FDA researchers and reviewers to use BCO, a writer, viewer, and editor was integrated into the CBER High-performance Integrated Virtual Environment (HIVE) platform. The display tool allows a user to browse the information in the BCO without needing a comprehensive understanding of the standard. The data is displayed hierarchically in a web interface as a series of expandable links. This allows an FDA reviewer or researcher to load a BCO they have received into HIVE and review how an analysis was performed in detail. Two BCOs can also be compared side by side with the differences highlighted. This can accelerate the review if a reviewer needs to see what changes a sponsor might have made to a bioinformatics pipeline between studies.

The writer tool creates a new BCO from the base BioCompute specification. A blank BCO can be created, but if a computation is user-selected, HIVE will automatically populate the parametric domain with the details of the computation. If multiple computations were used as a pipeline, the tool provides an easy way to add additional steps. HIVE can display a JSON of any computation that has been run in the platform, and that JSON can be copied and incorporated into the BCO within the creation tool. Finally, the user can fill out the description, the usability domain, and any additional domains and digitally sign the BCO. The editor tool allows a researcher to load and modify a BCO, exporting a new digitally signed copy when the BCO is finished.

### Galaxy BCO

The BioCompute Galaxy instance (https://galaxy.aws.biochemistry.gwu.edu/) is an open addition to the core Galaxy code and is developed by the BioCompute Team to better support data-intensive research. Users are able to assemble their pipeline, designate the outputs in the module boxes, and record the entire pipeline as a BCO made publicly available on Amazon Web Services (AWS).

### The CGC-integrated tool

The Cancer Genomics Cloud (CGC) [11] offers a web application for generating BioCompute Objects (CGC BCO App). The application provides an interactive web interface to help users browse, compose, and validate BCOs. To fulfill different types of use cases, the application allows the users to create and edit BCOs from user’s text input, from a workflow described in Common Workflow Language (CWL) [12] on the CGC, or from an existing task execution information on the CGC. CGC task information includes a workflow written in CWL as well as all workflow inputs and outputs, which we expect will reduce the amount of time required to generate a BCO. The CGC BCO app seamlessly integrates with the CGC via the CGC API, thus offering the capabilities to retrieve existing workflow and execution data from the CGC and to save BCOs to the cloud platform. This design strikes a balance between standardization and customization. The CGC BCO app sufficiently automates the process of BCO generation and largely reduces the amount of additional work to document data analysis pipelines without losing the flexibility to finetune the BCO details. Users can optionally save the generated BCOs to local machines as a JSON or PDF file, or push it to GitHub repositories for persistent storage.

The application was built with the R web framework Shiny [13], leveraging two R packages: tidycwl [14] and biocompute [15] to implement the essential support for working with the CWL and BCO specifications. During the PrecisionFDA BioCompute challenge, the application was hosted within a public CGC project and offered to student participants to run in the RStudio environment on the CGC. For advanced users, the application is containerized and offers flexible deployment options. The source code of the application is freely available from https://github.com/sbg/bco-app.

Grant information: This work was supported in part by the NIH through the National Cancer Institute for the Cancer Genomics Cloud. The Cancer Genomics Cloud, powered by Seven Bridges, is a component of the NCI Cancer Research Data Commons (datacommons.cancer.gov) and has been funded in whole or in part with Federal funds from the National Cancer Institute, National Institutes of Health, Department of Health and Human Services, under Contract No. HHSN261201400008C. and ID/IQ Agreement No. 17×146 under Contract No. HHSN261201500003I. This work was also supported by funding provided by the NIH to the PDXNet Data Commons and Coordination Center (NCI U24-CA224067).

